# PGC-1α regulates mitochondrial calcium homeostasis, SR stress and cell death to mitigate skeletal muscle aging

**DOI:** 10.1101/451229

**Authors:** Jonathan F. Gill, Julien Delezie, Gesa Santos, Shawn McGuirk, Svenia Schnyder, Stephan Frank, Martin Rausch, Julie St-Pierre, Christoph Handschin

**Author notes:** Correspondence to: Christoph Handschin, Biozentrum, University of Basel, Klingelbergstrasse 50/70, CH-4056 Basel, Switzerland. Present affiliation: Department of Biochemistry, Microbiology, and Immunology, University of Ottawa, K1H 8L1 Ottawa, Ontario, Canada (Julie St-Pierre).

## Abstract

Age-related impairment of muscle function severely affects the health of an increasing elderly population. While causality and the underlying mechanisms remain poorly understood, exercise is an efficient intervention to blunt these aging effects. We thus investigated the role of the peroxisome proliferator-activated receptor γ coactivator 1α (PGC-1α), a potent regulator of mitochondrial function and exercise adaptation, in skeletal muscle during aging. We demonstrate that PGC-1α overexpression improves mitochondrial dynamics and calcium buffering in an estrogen-related receptor α (ERRα)-dependent manner. Moreover, we show that sarcoplasmic reticulum stress is attenuated by PGC-1α. As a result, PGC-1α prevents tubular aggregate formation and fiber apoptosis in old muscle. Similarly, the pro-apoptotic effects of ceramide and thapsigargin were blunted by PGC-1α in muscle cells. Accordingly, mice with muscle-specific gain- and loss-of-function of PGC-1α exhibit a delayed and premature aging phenotype, respectively. Together, our data reveal a key protective effect of PGC-1α on muscle function and overall health span in aging.

**Statement of significance:** The loss of muscle function in aging results in a massive impairment in life quality, e.g. by reducing motor function, strength, endurance, the ability to perform daily tasks or social interactions. Unfortunately, the mechanistic aspects underlying age-related muscle disorders remain poorly understood and treatments improving the disease are extremely limited. We now show that PGC-1α, a transcriptional coactivator, is a key regulator of mitochondrial calcium homeostasis, cellular stress and death, all of which are linked to muscle aging and dysfunction. As a result, inhibition of the age-related decline in muscle PGC-1α considerably reduces aging of muscle and constitutes a promising target to prevent and treat the deterioration of muscle function in the elderly.

**Abbreviations:** BNIP3, BCL2/Adenovirus E1B 19kDa interacting protein 3; Cpt1b, carnitine palmitoyltransferase 1B; CSQ1, calsequestrin 1; Drp1, dynamin-related protein 1; ER stress, endoplasmic reticulum stress; ERRα, estrogen-related receptor α; Fis1, fission 1; GRP75, Glucose-Regulated Protein 75; IGFBP5, insulin like growth factor binding protein 5; IP3, inositol 1,4,5-trisphosphate; IP3R1, inositol 1,4,5-trisphosphate receptor type 1; Letm1, leucine zipper and EF-hand containing transmembrane protein 1; MAMs, mitochondria-associated ER membranes; Mcad, medium-chain acyl-CoA dehydrogenase; Opa1, optic atrophy 1; OXPHOS, oxidative phosphorylation; PGC-1α, peroxisome proliferator-activated receptor γ coactivator 1α; pH2AX, phospho-H2A Histone Family Member X; ppRB, phospho-prepro-retinoblastoma-associated protein; Puma, BCL2 Binding Component 3; ROS, reactive oxygen species; SR, sarcoplasmic reticulum; TA, tibialis anterior; TBP, TATA binding protein; TPG, thapsigargin; Ucp3, uncoupling protein 3; VDAC, voltage-dependent anion channel; XBP1, X-Box Binding Protein 1; Xiap, X-linked inhibitor of apoptosis protein

## Introduction

Muscle strength and mass progressively decline with age, leading to physical disability, and ultimately higher morbidity and mortality (1, 2). Although the cause of muscle aging is multifactorial (3), reduced mitochondrial function is a commonly observed phenomenon associated with muscle deterioration during aging (4, 5). For example, reduced mitochondrial biogenesis (6), decreased mitochondrial mass (7) and aberrant fission to fusion rates (8) concomitant with a mitochondrial turnover drop (9) have been reported in old muscle. In addition, increased mitochondrial DNA damage, depolarized and swollen mitochondria (4), diminished rates of oxidative phosphorylation (OXPHOS) as well as Krebs cycle activity (4, 7, 10) have been found in this context. Collectively, the ensuing reduced ATP production (11), impaired calcium homeostasis (12), elevated apoptosis (13) and increased levels of reactive oxygen species (ROS) (14) all contribute to age-associated muscle dysfunction.

Importantly, mitochondria, together with the sarcoplasmic reticulum (SR), control cellular calcium homeostasis (15, 16). These two organelles communicate via contact sites, termed mitochondria-associated ER membranes (MAMs), focal hotspots for the exchange of calcium as well as the synthesis and transfer of phospholipids, initiation of mitochondrial fission, mitophagy and signal transduction events (17). Aging impairs mitochondria-SR association and mitochondrial calcium uptake (18, 19), which can contribute to a dysregulation of cellular calcium homeostasis (20). Furthermore, all three structures, mitochondria, MAMs and the SR, have been linked to the initiation of cell death (21, 22) and increased apoptosis levels in old muscle (23–25). For example, apoptosis can be mediated in mitochondria via caspase-dependent and -independent pathways (26), both of which are rising with age (24, 26, 27). In addition, mitochondrial and nuclear DNA damage due to increased ROS levels also contribute to age-related muscle apoptosis (14). Finally, endoplasmic reticulum (ER) stress is also a strong promoter of apoptotic events (28).

Important regulators of mitochondrial biogenesis and function, most notably the peroxisome proliferator-activated receptor γ coactivator 1α (PGC-1α), are reduced in skeletal muscle in the aging process and have been associated with the decrease in mitochondrial function (29). Importantly, this deterioration can partly be restored by exercise (10), a strong promoter of PGC-1α transcription and activity (30). In addition, transgenic overexpression of PGC-1α extends lifespan in *Drosophila melanogaster* (31). Using gain- and loss-of-function mouse models for muscle PGC-1α, we aimed at providing a unifying and comprehensive study determining the role of PGC-1α in muscle aging, in particular in regards to calcium homeostasis and cell death.

## Results

### PGC-1α modulates mitochondrial dynamics and SR association in the aging muscle

In light of the strong decline in mitochondrial function in aging, we first performed a broad comparative analysis of young and old wild-type (WT), PGC-1α muscle-specific transgenic (mTg-PGC-1α) and PGC-1α muscle-specific knockout (mKO-PGC-1α) mice. In these skeletal muscle-specific gain- and loss-of-function models for PGC-1α, many of these parameters have been extensively assessed and reported in young animals (32–34), but data in old mice are scarce. Mitochondrial fission and fusion events are essential for proper mitochondrial physiology, but these programs controlling mitochondrial dynamics decline in aging (35). In skeletal muscle of 24 months old WT mice, we observed, along with a severe reduction of the *Pgc-1α* transcript (Supplemental Fig. S1a), an age-dependent decreased expression of the fusion genes *Mitofusin 1 and 2*(*Mfn1* and *2*) and *Optic atrophy 1 and 2* (*Opa1* and *2*) (Fig. 1a). Furthermore, muscle OXPHOS protein content as well as mitochondrial ATP generation and respiration were diminished in 24 months old WT mice (Supplemental Fig. S1b and c), in association with reduced spontaneous locomotion and endurance performance (Supplemental Fig. S1d and e). Intriguingly, loss of PGC-1α in muscles of young mKO-PGC-1α animals decreased OXPHOS protein (Supplemental Fig. S1b) and *Mfn1* gene expression levels (Fig. 1a) to values comparable to those obtained from aged WT muscles. Conversely, PGC-1α overexpression in young mTg-PGC-1α muscles strikingly up-regulated the levels of *Mfn1-2* and *Opa1-2*, as well as OXPHOS proteins, and prevented the age-linked reduction of these genes and proteins, as well as of mitochondrial respiration at 24 months (Fig. 1a; Supplemental Fig. S1b-c). In addition, muscle PGC-1α overexpression increased mtDNA amount as well as both mitochondrial density and size (Supplemental Fig. S2a-c), correlating with elevated expression of the estrogen-related receptor α (*Esrra*, ERRα) and the mitochondrial transcription factor A (*Tfam),* essential regulators of mitochondrial biogenesis and function (Supplemental Fig. S2d).

**Fig 1.**
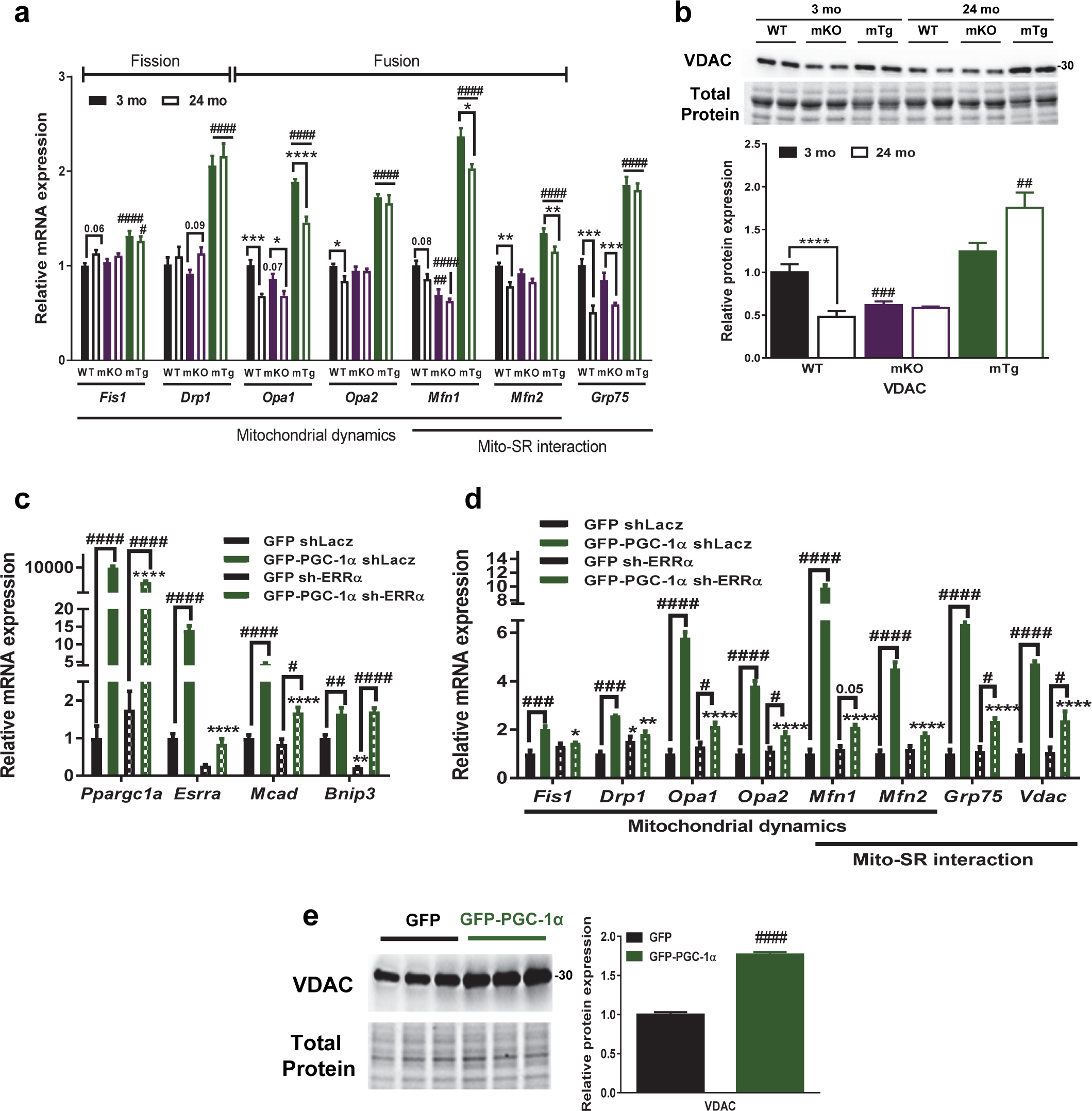
PGC-1α increases mitochondrial dynamics and SR association in old muscle and C2C12 cells in an ERRα-dependent manner. **(a)** Relative muscle mRNA levels of genes related to mitochondrial dynamics and SR-association (n=6). **(b)** Relative muscle VDAC protein levels (n=6) **(c and d)** Relative C2C12 cell mRNA levels of PGC-1α and ERRα targets and of genes related to mitochondrial dynamics and SR-association (n=3 independent experiments with 3 technical replicates). **(e)** Relative C2C12 cell VDAC protein levels (n=3 independent experiments with 3 technical replicates). Values are mean ± SEM. *P < 0.05; **P < 0.01; ***; P < 0.001; ****p<0.0001 indicate statistically significant differences between young and old animals of the same genotype or between cells with endogenous and overexpressed ERRα levels, # p<0.05; ## p<0.01; ### p<0.001; #### p<0.0001 indicate statistically significant differences between genotypes of age-matched animals or between cells with endogenous and overexpressed PGC-1α levels.

Besides their role in mitochondrial dynamics, mitofusin proteins, in particular Mfn2, constitute essential elements of MAMs, linking the SR and the mitochondrial compartments. In parallel with diminished *Mfn1* and *2* expression upon aging in WT muscles, a significant reduction of other essential mediators of the mitochondrial-SR communication, such as the glucose-regulated protein 75 (GRP75) and the voltage-dependent anion channel (VDAC) were detected (Fig. 1a and b). Interestingly, loss of PGC-1α in young mKO-PGC-1α muscles also decreased VDAC protein expression, while PGC-1α overexpression in mTg-PGC-1α animals resulted in higher levels of GRP75 gene and VDAC protein expression independent of age. We further demonstrated that overexpression of PGC-1α in C2C12 cells induces the expression of *Mfn1* and *2*, *Drp1*, *Fission 1* (*Fis1*), *Opa1*-*2*, *Grp75* and VDAC in an ERRα-dependent manner (Fig. 1c and d). Other PGC-1α target genes, for example the BCL2/Adenovirus E1B 19 kDa Interacting Protein 3 (*Bnip3*), were not affected by ERRα knockdown, attesting to the specificity of the PGC-1α-ERRα interaction in gene regulation (Fig. 1c). Together, these results clearly show that PGC-1α is not only an essential regulator of mitochondrial function and dynamics in young and old animals, but also prevents the age-associated decline of these systems.

### PGC-1α improves mitochondrial calcium handling during aging

A disrupted mitochondrial network and specifically the mitochondrial-SR association affects cellular calcium homeostasis (18, 19). Our group has previously demonstrated that PGC-1α modulates SR-controlled calcium levels in skeletal muscle (36), but the involvement of PGC-1α in regulating mitochondrial calcium uptake capacity and the exchange between mitochondria and the SR is unknown. We observed that aging significantly reduced the expression of genes involved in mitochondrial calcium uptake and transfer from the SR, including the leucine zipper and EF-hand containing transmembrane protein 1 (*Letm1*) and the inositol 1,4,5-trisphosphate (IP3) receptor type1 (*Itpr1*) in WT muscles (Fig. 2a). Interestingly, PGC-1α upregulation prevented the age decline of *Iptr1* and *Letm1* and significantly up-regulated genes related to mitochondrial calcium buffering in the muscle of old mice, including the mitochondrial calcium uniporter (*Mcu*) gene (Fig. 2a). The effect of PGC-1α on *Mcu* and *Letm1* was recapitulated in cultured myotubes (Fig. 2b). The PGC-1α-mediated induction of the expression of these two genes was furthermore dependent on ERRα (Fig. 2b). Intriguingly, the genes encoding proteins related to calcium exchange that form an organelle-spanning complex to control mitochondrial-SR interactions at MAMs were all downregulated in old WT mice, and rescued in old mTg-PGC-1α animals, including *Itpr1* on the SR side, *Grp75* providing a link between SR and mitochondria, and *Vdac* on the mitochondrial side. The tethering of this complex to both organelles is supported by the additional link provided by the mitofusins, which exhibit a similar regulation. Of note, this age-related dysregulation was at least in part exacerbated by knockout of PGC-1α. To test whether these changes affect calcium handling, mitochondrial calcium uptake in flexor digitorum brevis (FDB) muscle fibers isolated from old mTg-PGC-1α and WT mice were assessed. In line with the increased expression of MAM calcium exchange genes, mitochondria in FDB fibers of old muscle buffered more calcium when comparing mTG-PGC-1α to WT mice (Fig. 2c). Calcium removal kinetics were however not different between the genotypes (Fig. 2c). Next, the autonomous calcium buffering capacity of isolated mitochondria was investigated. We found that after the addition of 150 µM of calcium, calcium uptake was higher in mTg-PGC-1α compared to WT mitochondria (Fig. 2d and 2e). Since equal amounts of isolated mitochondria (275 µg) were used for these measurements, the results indicate that when normalized to mitochondrial amount, mitochondria from mTg-PGC-1α animals have an improved intrinsic calcium buffering capacity compared to mitochondria from WT mice. Thus, when considering the higher total mitochondrial mass in the gain-of-function model (Fig. 2f), the total mitochondrial calcium buffering capacity of skeletal muscle in mTgPGC-1α animals is even further amplified (Fig. 2g).

**Fig 2.**
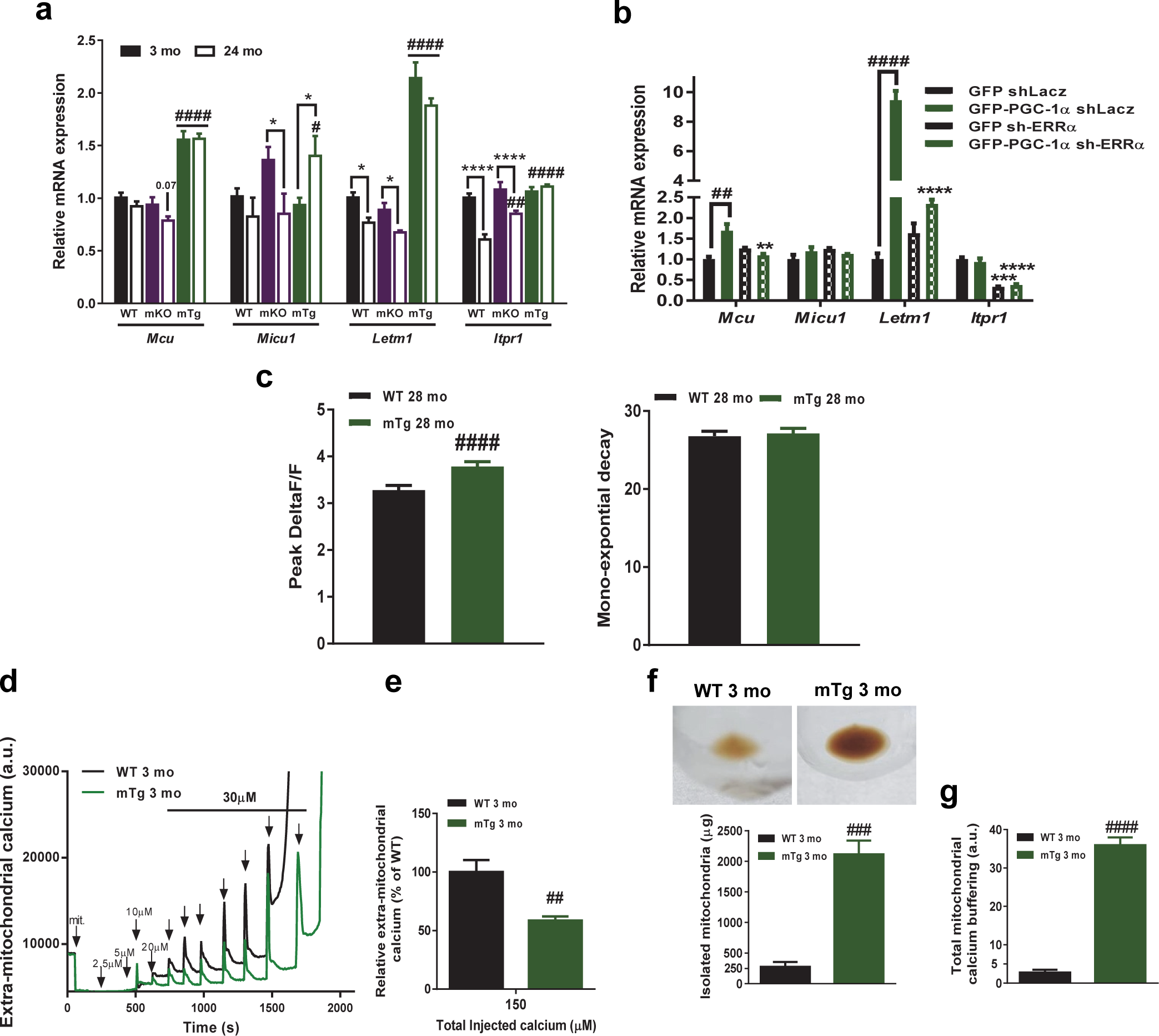
PGC-1α ameliorates mitochondrial calcium buffering. **(a)** Relative muscle mRNA levels of mitochondrial calcium buffering genes (n=6). **(b)** Relative C2C12 myoblasts mRNA levels of mitochondrial calcium buffering genes (n=3 independent experiments with 3 technical replicates). **(c)** Relative mitochondrial calcium uptake (left panel) and removal (right panel) in FDB fibers of 28 month-old animals **(d and e)** Representative extra-mitochondrial calcium traces upon calcium injection and quantification of relative extra-mitochondrial calcium levels after a total addition of 150 μm of calcium (n=4; 2 independent experiments with 2 muscles of each genotype used in each experiments). **(f)** Representative pictures of mitochondrial pellets after mitochondria isolation from hindlimb muscles and measure of isolated mitochondrial quantity. **(g)** Total mitochondrial calcium buffering capacity calculated by multiplying total mitochondrial quantity by the percentage of calcium imported in the mitochondria. Values are mean ± SEM. *P < 0.05; **P < 0.01; ***; P < 0.001; ****p<0.0001 indicate statistically significant differences between young and old animals of the same genotype or between cells with endogenous and overexpressed ERRα levels, # p<0.05; ## p<0.01; ### p<0.001; #### p<0.0001 indicate statistically significant differences between genotypes of age-matched animals or between cells with endogenous and overexpressed PGC-1α levels.

### PGC-1α prevents ER stress and tubular aggregate formation in the aging muscle

Dysregulated calcium exchange between the ER and mitochondria is linked to ER stress (37). To test whether the modulation in mitochondrial-SR interaction and mitochondrial calcium buffering affects the SR, the extent of ER stress was investigated in the muscles of our mouse cohorts. We observed that ER stress increases with age as demonstrated by elevated expression of the X-Box Binding Protein 1 (Xbp1) and the chaperon protein BIP in WT and mKO-PGC-1α muscles, but not to the same extent in mTg-PGC-1α animals (Fig. 3a and b). Furthermore, PGC-1α muscle deletion led to an increased activation of the calcium stress marker caspase 12, which was exacerbated with age (Fig. 3b). Consistently, poly-ubiquitination of proteins, indicative of proteasomal degradation of potentially misfolded proteins, was dramatically increased in mKO-PGC-1α muscles with age and reduced in both young and old muscles of mTg-PGC-1α mice (Supplemental Fig. S3a). Of note, the *in vivo* reduction of *Xbp1* mRNA levels and protein poly-ubiquitination by PGC-1α were recapitulated in differentiated C2C12 cells (Supplemental Fig. S3b and S3c).

**Fig 3.**
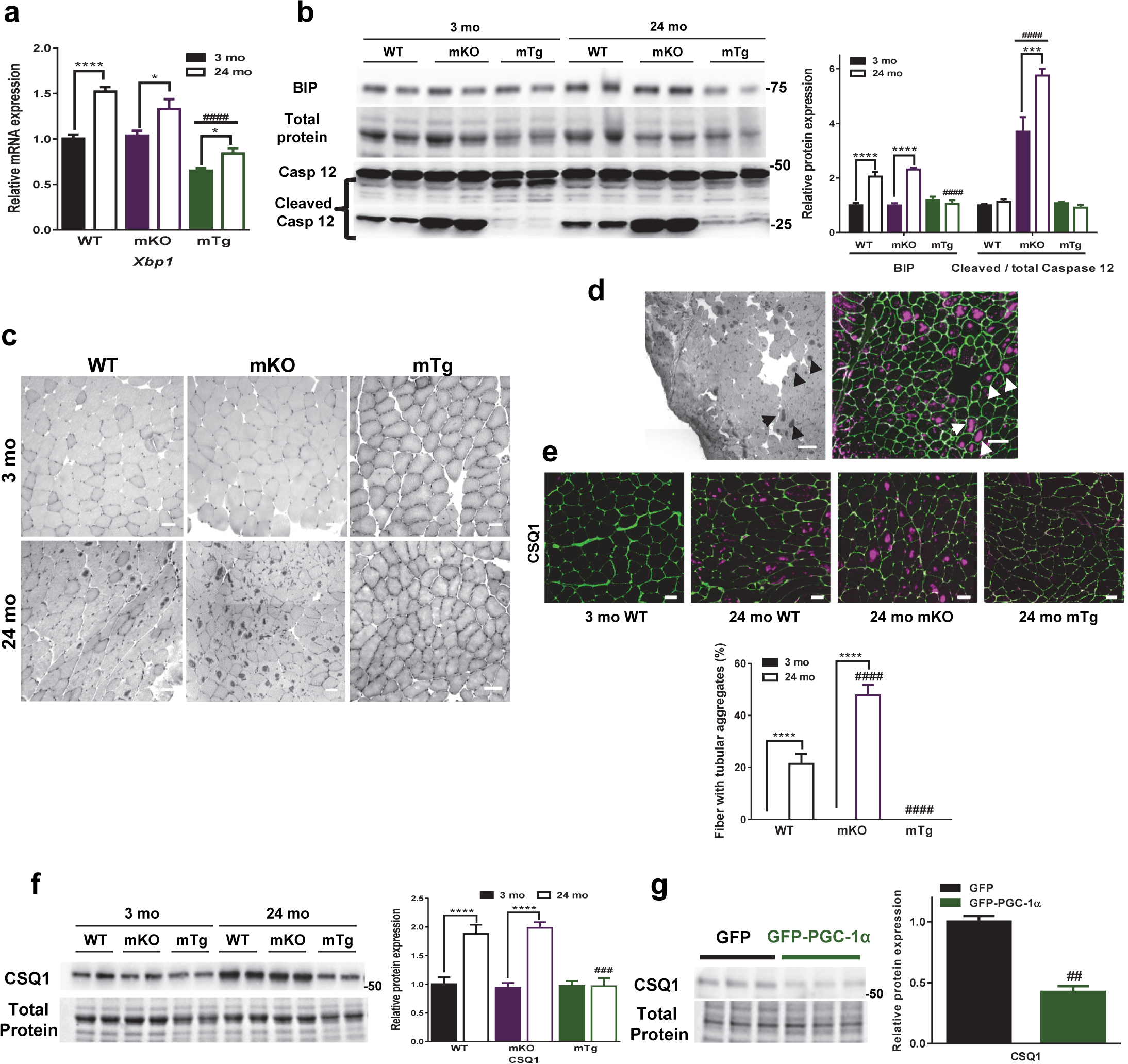
PGC-1α reduces ER stress during aging and prevents age-related tubular aggregate formation. **(a and b)** Relative muscle mRNA and protein levels of ER stress related genes (n=5-6). **(c)** Representative pictures of H&E stained tibialis anterior cryosection, scale bars represent 50 µm. **(d)** Colocalization of H&E labelled aggregates (left picture) with calsequestrin 1 staining (right picture) indicated by arrows, scale bars represent 100 µm. **(e)** Representative pictures of tubular aggregates stained with calsequestrin 1 antibody and quantification of the percentage of fibers containing tubular aggregates, scale bars represent 50 µm (n=6). **(f and g)** Calsequestrin 1 protein levels in muscles of young and old animals and in C2C12 myotubes (n=3 independent experiments with 3 technical replicates). Values are mean ± SEM. *P < 0.05; **P < 0.01; ***; P < 0.001; ****p<0.0001 indicate statistically significant differences between young and old animals of the same genotype, # p<0.05; ## p<0.01; ### p<0.001; #### p<0.0001 indicate statistically significant differences between genotypes of age-matched animals or between cells with endogenous and overexpressed PGC-1α levels.

Overload of SR function results in a compensatory increase in SR membranes and ultimately the development of tubular aggregates, e.g. as reported in muscle fibers of old mice (38). On histological sections, we noticed the presence of abnormal eosin-labeled structures in tibialis anterior (TA) muscles of old WT and mKO-PGC-1α animals (Fig. 3c), confirmed as tubular aggregates by positive staining for the SR marker calsequestrin 1 (CSQ1) in TA (Fig. 3d), and electron microscopy-based analysis of old WT and mKO-PGC-1α gastrocnemius muscles (Supplemental Fig. S4a). Interestingly, the age-dependent occurrence of tubular aggregates doubled in TA muscles of old mKO-PGC-1α compared age-matched WT animals (Fig. 3e). Remarkably, no tubular aggregates were detected in fast and mixed muscles of aged mTg-PGC-1α mice (Fig. 3c and e and Supplemental S4a). Tubular aggregates were likewise reduced in the oxidative soleus muscle (Supplemental Fig. S4b) and succinate dehydrogenase (SDH)-positive fibers of other muscle beds in old WT and mKO-PGC-1α mice (Supplemental Fig. S4c), suggesting a link to the respective metabolic fiber type. The formation of tubular aggregates is closely associated with the accumulation of CSQ1 and other SR proteins, which might subsequently overwhelm the unfolded protein response (39, 40). Thus, along with the absence of tubular aggregate formation in muscle of old mTg-PGC-1α mice, elevation of muscle PGC-1α levels prevented the age-related increase of CSQ1 protein levels that is observed in WT and mKO-PGC-1α mice (Fig. 3f). The acute reduction of CSQ1 protein and transcript levels in cultured myotubes overexpressing PGC-1α implies that this effect is at least in part directly linked to PGC-1α, and not only relies on indirect, secondary effects of a fiber type shift (Fig. 3g). Of note, in addition to the tubular aggregates, electron microscopy also revealed other age-linked, abnormal structures of unknown origin and function in muscles fibers of old mKO-PGC-1α and WT mice that were absent from the young animals of all three genotypes and from old mTg-PGC-1α mice (Supplemental Fig. S4d) indicating further age-associated abnormalities and damage in skeletal muscle that can be prevented by high levels of PGC-1α.

### PGC-1α alleviates muscle cell death during aging

Abnormal mitochondrial function, ER stress response, mitochondrial-SR interaction and calcium homeostasis have all been described in the context of the initiation of cell death (22, 41–43). We therefore assessed whether age-associated apoptosis in skeletal muscle was affected by modulation of PGC-1α. P53, a major regulator of cell death induction upon DNA damage and other stress conditions, was upregulated during aging of WT and mKO-PGC-1α muscles, but not in mTg-PGC-1α muscle overexpressing PGC-1α (Fig. 4a). Compared to aged WT animals, muscles of old mKO-PGC-1α and mTg-PGC-1α mice exhibited, respectively, trends toward a 2-fold increase (p=0.06) and a 2-fold decrease (p=0.05) in P53 protein levels. Interestingly, mRNA levels of the pro-apoptotic insulin-like growth factor binding protein 5 (*Igfbp5*) and of the p53 targets *p21* and BCL2-binding component 3 (*Puma)* followed a similar pattern of expression, implying increased activity of P53 in old WT and mKO-PGC-1α, but not mTg-PGC-1α mice (Fig. 4b). The transcript levels of the cell survival-related gene X-linked inhibitor of apoptosis protein (*Xiap*) was down-regulated with age in WT and mKO-PGC-1α muscles and cyclin D transcript (*Ccnd1*) expression was lower in old mKO-PGC-1α muscles relative to age-matched WT muscles. PGC-1α significantly elevated the expression of all pro-survival genes in young, and *Xiap* mRNA levels in old, muscles of mTg-PGC-1α mice (Fig. 4b). Caspase 3 cleavage, a marker of cell death, was increased with age in mKO-PGC-1α mice and significantly higher in muscles of old mKO-PGC-1α mice relative to muscles of age-matched WT animals (Fig. 4c). Additionally, in old muscle tissues, the smallest cleavage product of caspase 3 was reduced by PGC-1α overexpression. Taken together, our findings suggest a protective function of muscle PGC-1α against age-induced muscle cell death.

**Fig 4.**
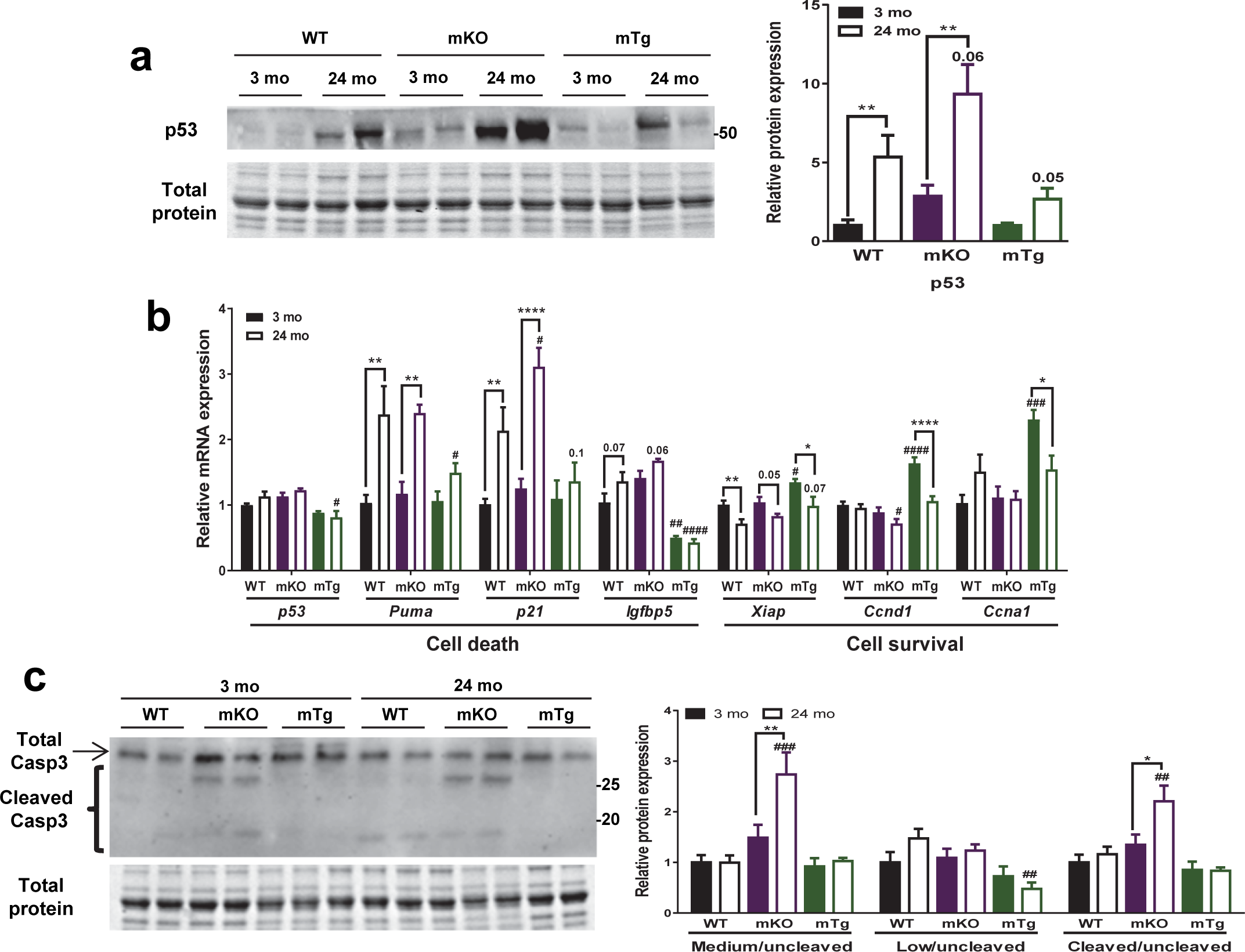
PGC-1α inhibits age-related muscle cell death. **(a-c)** Relative muscle mRNA and protein levels of cell death and cell survival related genes (n=5-6). Values are mean ± SEM. *P < 0.05; **P < 0.01; ***; P < 0.001; ****p<0.0001 indicate statistically significant differences between young and old animals of the same genotype, # p<0.05; ## p<0.01; ### p<0.001; #### p<0.0001 indicate statistically significant differences between genotypes of age-matched animals.

### PGC-1α protects against ceramide- and thapsigargin-induced cell death

To evaluate the direct influence of PGC-1α on cell death, we used ceramide, which acts as a second messenger for apoptosis linked to ER stress, mitochondrial impairment and dysregulation of calcium homeostasis, hence a number of events that we also observed in muscle aging *in vivo* (44). We observed that PGC-1α overexpression protected C2C12 cells following ceramide exposure as evaluated by microscopy and by a propidium iodide-based cell death assay (Fig. 5a and b). Of note, similar findings have previously been reported in HeLa cells (45). Furthermore, PGC-1α overexpression in muscle cells abolished the induction of p53, reduced the cleavage of caspase 3 proteins and alleviated the increase in expression of the DNA damage marker phospho-H2A histone family member X (pH2AX) in response to ceramide treatment (Fig. 5c). Inversely, protein levels of the phosphorylated form of the phospho-prepro-retinoblastoma-associated protein (ppRb), which represents an inactive state of this protein involved in triggering apoptosis upon DNA damage (46), were increased by PGC-1α (Fig. 5c). Moreover, PGC-1α upregulation abrogated ceramide-dependent changes of gene expression related to cell death and survival (Fig. 5d). Next, we assessed whether PGC-1α protects myocytes from cell death specifically induced by cytosolic calcium dysregulation. Therefore, we treated myocytes with thapsigargin (TPG), a non-competitive inhibitor of the ATPase sarcoplasmic/endoplasmic reticulum Ca2+ transporting (SERCA), which blocks SR calcium uptake and promotes apoptosis through a dramatic increase of cytosolic calcium and ER stress (47). Similar to its protective role after ceramide treatment, PGC-1α protected muscle cells from TPG-induced apoptosis (Fig. 6a and b). This protective effect was accompanied by a reduction of TPG-mediated caspase 3 activation and pH2AX protein elevation (Fig. 6c). At the same time, the downregulation of ppRB protein levels by TPG was prevented by PGC-1α overexpression (Fig. 6c). Notably, unlike ceramide, TPG did not affect p53 protein levels, indicating that ceramide and TPG induce apoptosis through at least in part different pathways, both of which are affected by PGC-1α. Together, these data suggest an essential role for PGC-1α in the prevention of cell death.

**Fig 5.**
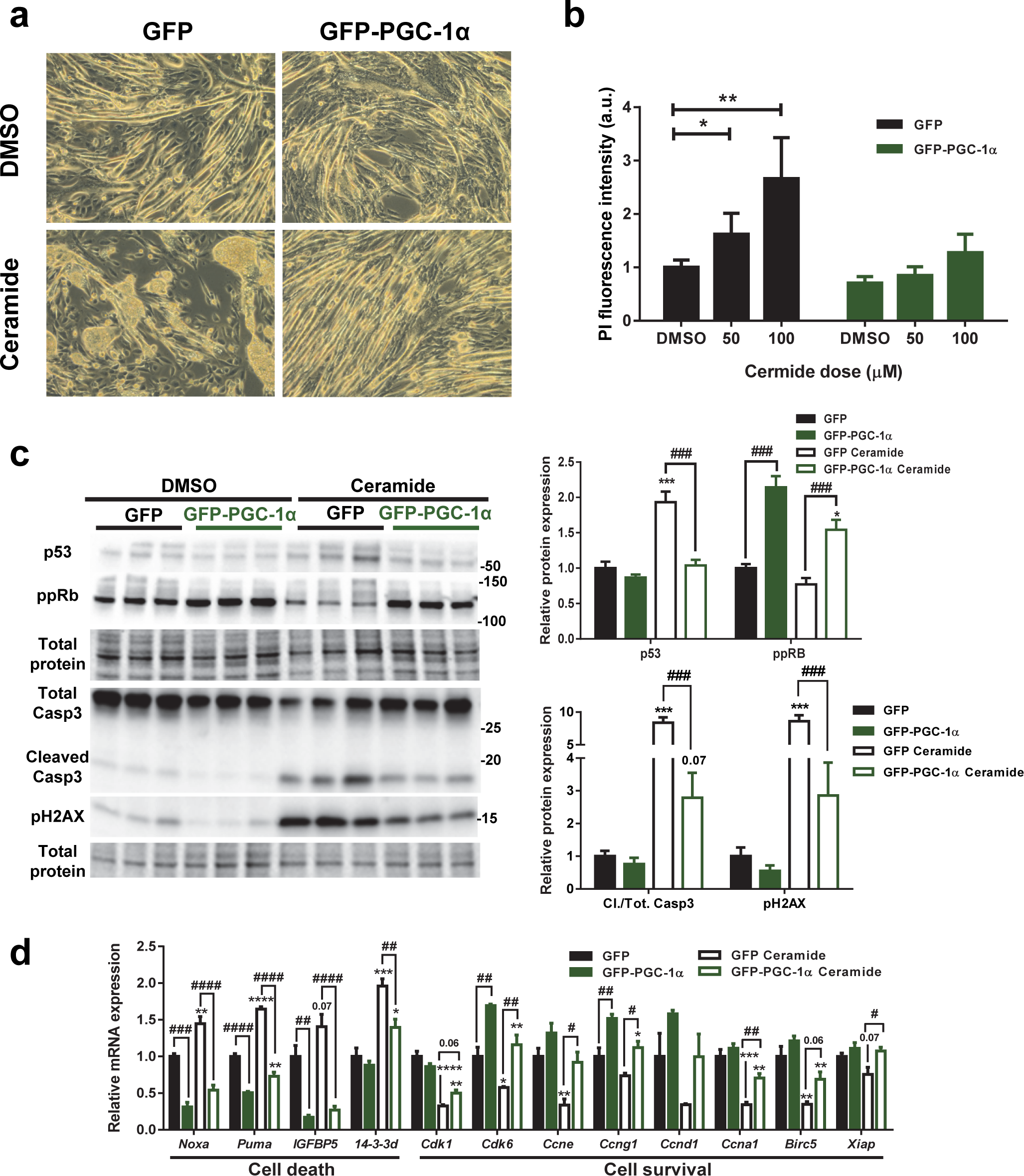
PGC-1α protects from ceramide-induced cell death. **(a and b)** Representative pictures of myotubes and propidium iodide incorporation in myoblasts with endogenous or increased PGC-1α levels after ceramide or DMSO treatment. **(c)** Relative protein levels of P53 and ppRB in myotubes and Caspase 3 and pH2AX in myoblasts. **(d)** Relative myotube mRNA levels of cell death and cell survival markers (n=3 independent experiments with 3 technical replicates). Values are mean ± SEM. *P < 0.05; **P < 0.01; ***; P < 0.001; ****p<0.0001 indicate statistically significant differences between cells treated with DMSO and ceramide, # p<0.05; ## p<0.01; ### p<0.001; #### p<0.0001 indicate statistically significant differences between cells with endogenous and overexpressed PGC-1α levels.

**Fig 6.**
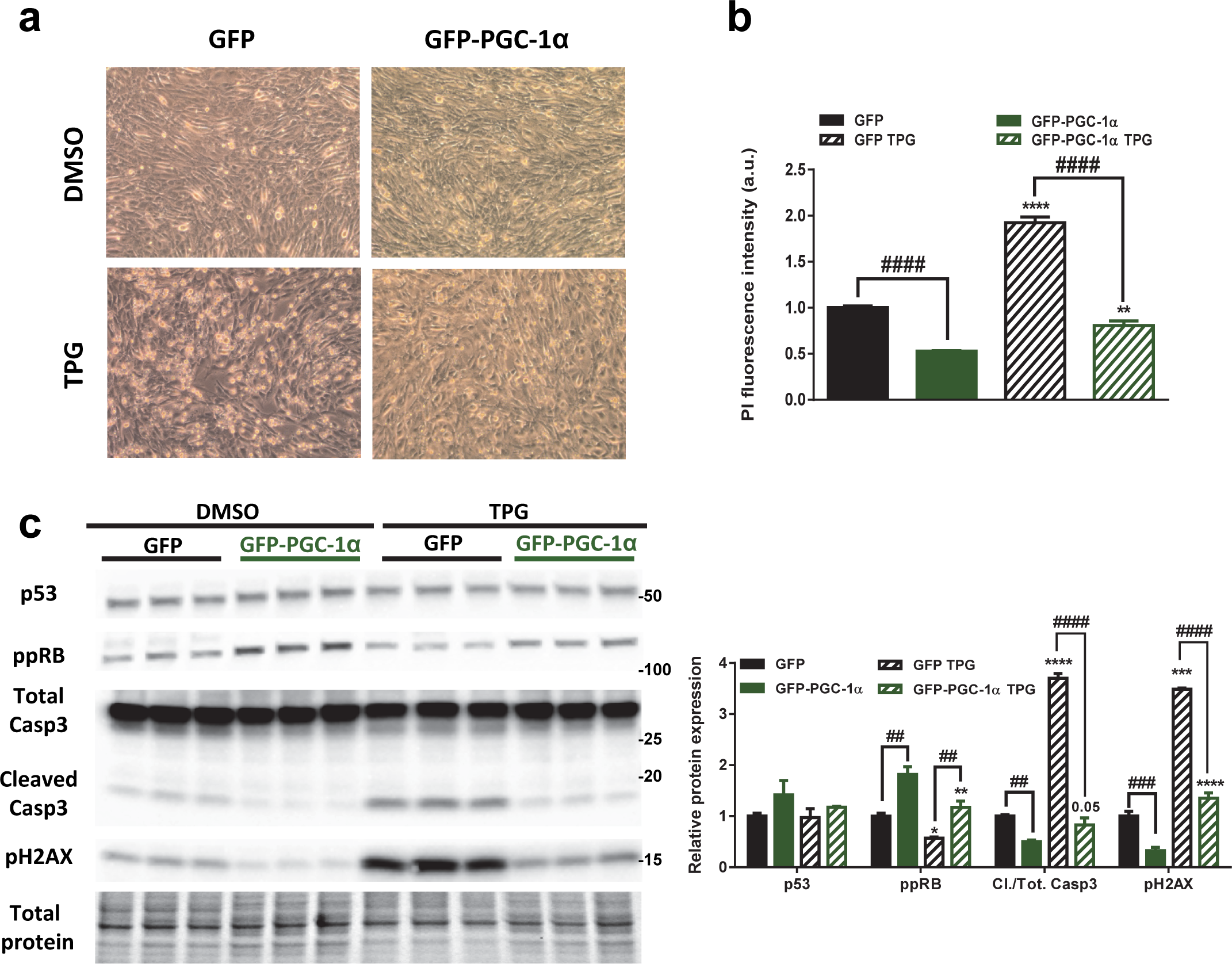
PGC-1α inhibits TPG-induced cell death. **(a and b)** Representative pictures and propidium iodide incorporation in myoblasts with endogenous or increased PGC-1α levels after TPG or DMSO treatment. **(c)** relative myoblast protein levels of cell death and cell survival markers after TPG or DMSO treatment. (n=3 independent experiments with 3-4 technical replicates). Values are mean ± SEM. *P < 0.05; **P < 0.01; ***; P < 0.001; ****p<0.0001 indicate statistically significant differences between cells treated with DMSO and TPG, # p<0.05; ## p<0.01; ### p<0.001; #### p<0.0001 indicate statistically significant differences between cells with endogenous and overexpressed PGC-1α levels.

## Discussion

PGC-1α is a key regulator of mitochondrial biogenesis, dynamics and function in young animals. Moreover, the effects of this transcriptional coactivator extend to other organelles and cell compartments, such as the unfolded protein response after exercise (48) and SR-controlled calcium homeostasis (36). We describe here a novel PGC-1α-regulated pathway involving the collective control of the function and interaction between mitochondria and the SR centered on calcium homeostasis. Our findings accordingly provide a mechanistic and functional link between the observed decline in PGC-1α expression and the ensuing reduction in mitochondrial dynamics and activity in muscle at old age that have been reported previously (4, 6, 10, 29).

Mitochondrial calcium uptake is important for electron transport chain function and hence ATP production, but also significantly contributes to the modulation of intramyocellular resting calcium levels (15). For example, the age-related decline in mitochondrial calcium buffering capacity leads to a rise in intracellular calcium levels in old muscle (18, 19, 49). The improved calcium buffering capacity of mitochondria from muscles overexpressing PGC-1α can therefore blunt much of the stress exerted by increased intracellular calcium in this context. Moreover, improved mitochondrial function and ATP production help to maintain calcium re-uptake into the SR via the ATP-dependent calcium pumps (50). Concurrently, the reduction in poly-ubiquitinated proteins and the expression of Xbp1 and BIP indicate a PGC-1α-dependent alleviation of the SR burden. Complete abrogation of tubular aggregates formation by PGC-1α further illustrates the muscle protection from ER stress development and escalation to tubular aggregates during aging.

Tubular aggregates are predominantly composed of small, densely packed tubules arising from sarcoplasmic reticulum (39) and are associated with both natural and premature aging in mice (38, 51). In humans, tubular aggregates have likewise been reported in old muscle (52), but are also prominently observed in several other pathological conditions, including peripheral neuropathies, amyotrophic lateral sclerosis and myotonic dystrophy (53–55). Moreover, these abnormal structures represent the predominant symptom in tubular aggregate myopathy (55). Due to their calcium loading capacity (56), tubular aggregates are alternatively thought to be a compensatory mechanism to counteract age-associated cellular calcium increase, rather than a consequence of dysregulated calcium metabolism. Here, we report that PGC-1α overexpression fully protects muscle from tubular aggregate formation in old mice, while muscle ablation of the PGC-1α gene markedly exacerbates tubular aggregate occurrence during aging (Fig. 3 and Supplemental Fig. S4) or, as previously reported, upon denervation (57). Moreover, a link between impaired mitochondrial calcium handling and tubular aggregate formation was reported in young mitochondrial calcium uptake 1 (MICU1) knockout mice, which recapitulate some of the phenotype of old WT and young and old PGC-1α knockout muscle, including mitochondrial dysfunction, impaired muscle function and the formation of tubular aggregates (58). In parallel to improving calcium homeostasis and downregulation of proteins that accumulate in tubular aggregates such as CSQ1, a central role for PGC-1α in this context is further substantiated by our SDH staining and electron microscopy data as well as other studies that revealed a strong preference for tubular aggregate formation in glycolytic fibers (59, 60). The potent effect of PGC-1α in driving an oxidative muscle fiber shift (32, 33, 61) thus likely further contributes to the inhibition of tubular aggregate formation. In light of our data, it would be interesting to study whether increased muscle PGC-1α levels prevent tubular aggregate formation in mice with tubular aggregate-associated myopathies (51, 62).

Mitochondrial dysfunction, ATP depletion, calcium dysregulation and ER stress, as observed in old WT and mKO-PGC-1α mice, are strong promoters of cell death (37, 42, 63). The protective effect of muscle PGC-1α on mitochondrial and SR function could thus directly or indirectly result in the observed reduction of cell death initiation. Analogous to the results *in vivo*, the induction of cell death in C2C12 cells upon ER and mitochondrial stressors (e.g. as evoked by ceramide) or abnormal cellular calcium elevation (e.g. as triggered by TPG) is inhibited when PGC-1α is overexpressed. Interestingly, the anti-apoptotic function of PGC-1α has previously been suggested in a number of tissues, including the retina (64), vascular endothelial cells (65), neurons (66) and skeletal muscle (67). Our findings now provide direct evidence of a broader PGC-1α-controlled program that links functional mitochondria, SR and calcium handling to cell death regulation in muscle, which becomes compromised with decreased PGC-1α expression during aging, and ultimately leads to higher rates of myofiber apoptosis. The effect of PGC-1α on P53, pH2AX and ppRb is reminiscent of findings in vascular endothelial cells, in which a telomere-P53-PGC-1α axis has been postulated to be involved in regulating apoptosis and senescence (68, 69). A similar signaling cascade could accordingly be involved in modulating DNA damage and cellular senescence in old skeletal muscle.

In summary, our results demonstrate that PGC-1α, in close relation with ERRα, is central in the coordinated control of SR and mitochondrial calcium homeostasis, thereby preventing SR stress and cell death (Fig. 7). Intriguingly, this control is exerted in a context-specific manner: for example, while PGC-1α upregulates the unfolded protein response to cope with the acute consequences of exercise (48), increased muscle PGC-1α in old muscle alleviates the aging-related burden on the SR by lowering the ER stress response. Accordingly, mKO-PGC-1α mice in many regards exhibit a premature aging phenotype (present results and ref. (61)). Along the same line, these animals have previously been reported to suffer from exacerbated age-related glucose intolerance and systemic inflammation (70). This is different from cardiac tissue, where a reduction in PGC-1α gene expression has a limited effect on aging, mainly affecting mitochondrial gene expression (71). Inversely, PGC-1α elevation improved both cardiac (71) and skeletal muscle in old mTg-PGC-1α mice, overall positively affecting health-adjusted life expectancy (health span), thus the length of time of life in a healthy state, as evidenced by preserved locomotion and exercise capacity (Supplemental Fig. S1d and e), as well as balance and motor coordination (61). Our findings are furthermore corroborated by a recent publication reporting a delayed aging process of mTg-PGC-1α animals in terms of global gene expression patterns, markers for mitochondrial function and muscle wasting, as well as morphology of the neuromuscular junction (72). Interestingly, while mTg-PGC-1α exhibit such an extension in health span, alteration of muscle PGC-1α expression did not affect overall life span in our study (Supplemental Fig. S5). A modest increase (∼5%) in median life span has however been recently been reported in those mTgPGC-1α mice that reached a minimal age of 800 days, but not when all animals were analyzed (72), indicating a subtle beneficial effect of muscle PGC-1α on survival in old mice. Collectively, our present findings reveal a novel cascade of cellular processes explaining how PGC-1α affects muscle health in aging, involving SR-mitochondrial interaction linked to calcium handling, SR stress and cell death (Fig. 7). In the light of our present and other recent findings (61, 70, 72), PGC-1α modulation, e.g. by exercise or pharmacological interventions, represents an attractive approach to reduce weakness, frailty and other pathological alterations associated with skeletal muscle aging, along with additional potential benefits on the heart (71).

**Fig 7.**
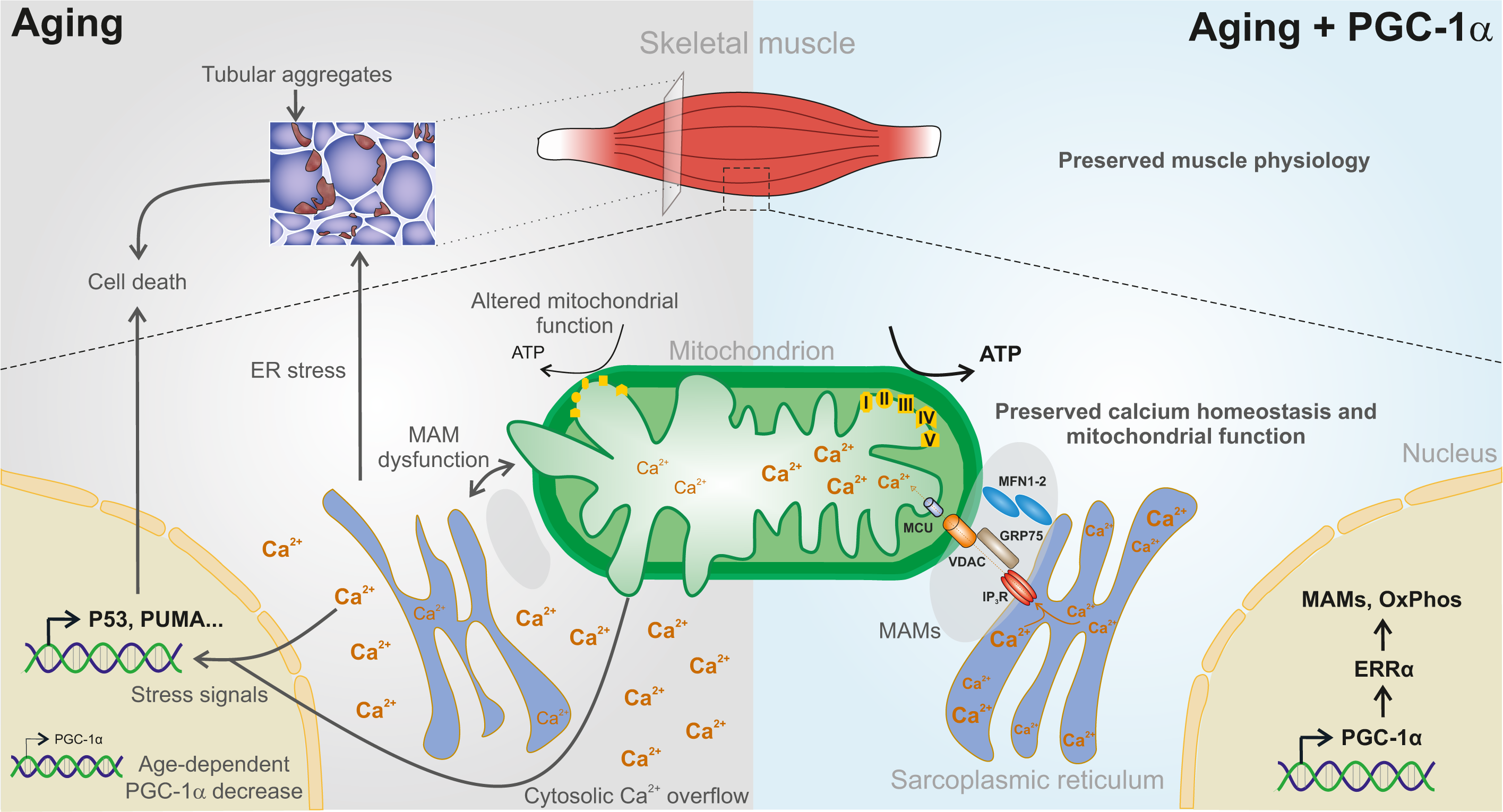
PGC-1α regulation of calcium and cell death during skeletal muscle aging. Aging and PGC-1α downregulation impair mitochondrial respiration, calcium import and SR-association leading to exacerbated cytosolic calcium increase and SR-stress, which ultimately results in tubular aggregate formation. Those cellular dysregulations all contribute to promote muscle apoptosis. PGC-1α elevation improves mitochondrial function and calcium buffering, preserving calcium homeostasis and preventing tubular aggregate development during aging. Together with the reduction of those cellular insults, PGC-1α inhibits apoptosis initiation, thereby further protecting muscle from cell death.

## Methods

### Animals and reagents

Mice with muscle specific PGC-1α deletion (mKO-PGC-1α) or overexpression (mTgPGC-1α) were previously described (32–34). C57/Bl6 wild-type (WT) mice were obtained from Janvier. Male mice were studied at 3 and 24 months of age unless otherwise stated. All experiments were performed in accordance with the federal guidelines for animal experimentation and were approved by the Kantonales Veterinäramt of the Kanton Basel-Stadt.

Primary antibodies were obtained from Cell Signaling, Thermo Scientific, Abcam, Sigma or Enzo, and secondary antibodies from Dako, Jackson Immunoresearch or Life Technology. General reagents were purchased from Sigma Aldrich.

### Mouse muscle preparation

Mice were sacrificed by CO2 inhalation. Muscles were harvested and either snap-frozen in liquid nitrogen for protein and RNA extraction, frozen in cooled-isopentane and embedded in tragacanth for cryosection staining, immediately used for determination of respiration or calcium uptake in isolated mitochondria or fixed for electron microcopy.

### Treadmill experiment and spontaneous locomotor activity

To assess locomotor performance, mice ran in an open treadmill (Columbus instruments). Mice were acclimatized to the treadmill for 5 min at 8 m/min followed by 5 min at 10 m/min, at an incline of 5° during two consecutive days. After one resting day, an exhaustion test was performed at an inclination of 5° and a starting running speed of 4.8 m/min which was then increased by 1.6 m/min every 3 min until a maximum speed of 29 m/min. Maximal running speed was recorded when exhaustion was reached. Blood lactate levels were measured from the tail vein before and 10 min after the endurance test with a lactate plus meter (Nova biomedical).

Spontaneous locomotor activity was recorded by counting number of beam breaks in an indirect calorimetric system (CLAMS, Columbus Instruments) in 15 min intervals. Data were analyzed after one day of acclimatization for 2 days.

### Histology

Tragacanth-embedded muscles were cut to 8 µm sections with a cryostat (Leica, CM1950). H&E staining was performed as described in the DMD_M.1.2.007 SOP (http://www.treat-nmd.eu/downloads). For SDH staining, sections were incubated in SDH buffer (phosphate buffer 0,035M, Na-succinate 0.1M, teranitro blue trezolium salt 0.1%, phenazin methosulfat 0.1%), rinsed with water, fixed with 4% formalin and rinsed with water again before being mounted with CC/Mount (Sigma, C9368). For calsequestrin 1 labelling, cryosections were fixed for 20 min with 4% PFA in PBS (Sigma-Aldrich, D8537) at room temperature. Sections were blocked in PBS supplemented with 0.4% Triton X-100 (Sigma-Aldrich, 93426), 3% Goat Serum (Sigma-Aldrich, G9023), 1% BSA (Sigma-Aldrich, A9418). Sections were blocked for 30 min at room temperature and incubated with primary and secondary antibodies for one hour at room temperature. 3 washes of 5 min in PBS were performed before and after antibody incubation. Primary and secondary antibodies were diluted in blocking solution. Calsequestrin 1 (Thermo Scientific, MA3-913) and laminin (Sigma, L9393) antibody dilutions were 1/250 and 1/5000 respectively. AlexaFluor 488 (Life Technology, A-11008) and 647 (A-21242) secondary antibody dilutions were 1/500 and 1/250 respectively. Labelled sections were mounted with ProLong Gold Antifade reagent (Life Technologies, P36931). Images of calsequestrin 1 immunolabellings and H&E stainings were taken at the imaging core facility of the Biozentrum with the FEI MORE microscope using a 20x or 40X magnification lens, keeping the same acquisition settings for all compared samples. Numbers of tubular aggregates were quantified in all fibers of each stained muscles using ImageJ software with original images. Representative images were however adjusted for brightness and contrast.

### Genomic DNA/RNA extraction and qPCR

- Genomic DNA For genomic DNA extraction, crushed gastrocnemius muscles were shaken overnight at 55° in 600 µl lysis buffer (10mM Tris-HCl, 1mM EDTA, 0.1% SDS 5% Proteinase). Lysates were centrifuged at 8000g for 15 min at room temperature and supernatants transferred to fresh tubes. Residual RNA was removed by incubation with RNase A (20mg/ml) for 30 min at 37°C under constant agitation. RNase A was inactivated for 10 min at 95°C and samples were cooled-down to room temperature. One volume of phenol/chloroform/isoamylalcohol 25:24:1 was added and samples were centrifuged at 8000g for 15 min at room temperature. The aqueous phase was transferred to a new tube before addition of one volume of chloroform. The centrifugation step was repeated and the new aqueous phase was transferred again to a new tube where one volume of isopropanol containing 0.3M of sodium acetate was added. Samples were gently mixed and placed at −20°C for 30 min. DNA was recovered by centrifugation at 8000g for 15 min at 4°C. DNA pellets were washed 2 times with 1ml of ice-cold 75% ethanol, dried during few minutes and finally resuspended in ddH2O. An amount of 0.1µg of gDNA was used for qPCR.
- RNA extraction and qPCR Total RNA was isolated from powdered muscles using lysing matrix tubes (MP Biomedicals 6913-500) and TRI Reagent (Sigma-Aldrich T9424). Total RNA was recovered from cells using the Direct-zol RNA MiniPrep kit (Zymo Research R2050 according to the manufacturer’s instructions). After treatment with RNase-free DNase (Invitrogen 18068-015), 1µg of RNA was used for reverse transcription using the SuperScript II reverse transcriptase (Invitrogen 18064-014). The level of relative genomic DNA or mRNA was quantified by real-time PCR on a Light Cycler 480 system (Roche Diagnostics) using FastStart essential DNA probe master mix (Roche Diagnostics 06402682001). Relative quantification of mRNA for gene expression comparison was performed with the ΔΔCT method using the TATA binding protein (TBP) gene as reference. The quantification of mitochondrial DNA copy number was done using the same method by normalizing the average of COX1 ATP6 and ND1 DNA copy number by the average copy number of the nuclear genes beta globin and 34B6. Beta globin and TBP levels were similar between genotypes in a given experimental condition. Primer sequences are listed in Supplemental Table S1.

### Protein extraction and Western Blot

Quadriceps muscles were crushed on dry ice and homogenized in 250μL of ice-cold lysis buffer (50 mM Tris-HCl (pH 7.5), 250 mM sucrose, 0.25% Nonidet P 40 substitute, 1 mM EDTA, 1 mM EGTA, 50 mM NaF, 5 mM Na 0.1% DTT, fresh protease, and phosphatase inhibitor mixture). Samples were then incubated for 30 min at 4 °C under constant agitation at 1,300 rpm, before being centrifuged at 13,000×g for 10 min at 4 °C. The resulting supernatants were transferred to a fresh tube and protein concentrations were measured by Bradford assay (Bio-Rad, 500-0205). Total proteins from C2C12 cells were extracted after an ice-cold PBS wash following the same procedure. Equal amounts of protein were separated on mini-TGX 4-20% stain free pre-cast gel (Biorad, 4568096). Proteins were labelled with trihalo compounds of the stain-free gel by exposing the gel to UV for 1 min. Gel and nitrocellulose membranes were equilibrated for 5 min in transfer buffer and proteins were transferred on the nitrocellulose membrane during 1h under a constant voltage of 100V. Membranes were then blocked 1 h at room temperature with 5% milk or BSA diluted in TBS-T and washed 2 times 5 min with TBS-T. Proteins of interest were then labelled overnight at 4°C with primary antibodies diluted in TBS-T containing 0.02% sodium azide and either 3% milk or 3% BSA. Membranes were washed three times 5 min with TBS-T. Membranes were then incubated 1 h at room temperature with peroxidase-conjugated secondary antibodies diluted in TBS-T containing 3% milk or BSA. Membranes were then washed 3 times for a total of 15 min. Antibody binding was revealed using the enhanced chemiluminescence HRP substrate detection kit SuperSignal™ West Dura Extended Duration Substrate (Thermoscientific, #34076) and imaged using a fusion FX imager. Proteins of interest were normalized against total protein content determined using the trihalo compounds labeling. Two reference samples were loaded in the different gels for inter-gel normalization. Quantification of proteins was done with fusion FX software. Blocking and antibody solutions, antibody information and dilutions are described for each protein of interest in Supplemental Table S2. All primary antibodies were diluted at 1/1000 except Calsequestrin 1 antibody that was diluted at 1/3000. All secondary antibodies were diluted at 1/10000.

### C2C12 growth, differentiation and viral infection

C2C12 cells were cultured DMEM (Sigma-Aldrich D 5796), supplemented with 10% fetal bovine serum (HyClone Laboratories, Inc., Logan, UT), 4.5 mg/ml glucose and 1% penicillin/streptomycin and were incubated at 37 °C with 5% CO2. Myotube differentiation was achieved by incubation of 95% confluent myoblasts in differentiation medium (DMEM with 2% horse serum, 4.5 mg/ml glucose and 1% penicillin/streptomycin) during 4 days. PGC-1α overexpression was performed with adenoviral vectors expressing bicistronic GFP-PGC-1α or GFP alone as control. Infection was initiated in 50% confluent myoblasts or in myotubes after 4 days of differentiation. ERRα knock-down combined with PGC-1α overexpression was performed in C2C12 myoblasts by infecting cells with adenoviral vectors containing specific shRNA sequences against ERRα (shEsrra) or Lacz (shLacz) simultaneously with viruses used to study PGC-1α upregulation.

For cell death experiments, 2 days after infection, muscle cells were treated for 8 hours with 50 µM or 100 µM ceramide (Sigma-Aldrich 01912) or for 24h with 1µM thapsigargin (Sigma-Aldrich T9033). 0.1% DMSO (Sigma-Aldrich 276855) was used as control in both treatments. Pictures of myocytes after treatment were taken using Leica DMI4000B microscope with a 10x magnification. After myoblast exposure to ceramide or thapsigargin, cell death assays were performed by exchanging treatment medium with fresh medium containing 5 µM of propidium iodide. Cells were incubated for 30 min at 37°C with 5% CO2 and propidium iodide incorporation reflecting cell death was measured using a tecan infinite M1000 multiplate reader.

### Mitochondrial respiration assay

Quadriceps muscles were quickly rinsed in PBS and in PBS with 10 mM EDTA and finely minced in Petri dishes filled with 2ml of isolation buffer 1 (EDTA 10 mM, Dmannitol 215 mM, sucrose (0.075M), free-fatty acid BSA (Sigma-Aldrich) 0.1%, HEPES 2 mM pH 7.4 in distilled water). Muscle solutions were transferred in Potter-Elvehjem grinders for homogenization with manual pestle. Homogenates were centrifuged for 10 min at 700g. Supernatants were transferred and centrifuged for 10 min at 10500g. Pellets were resuspended in 500µl of isolation buffer 2 (EGTA 3 mM, D-mannitol 215 mM, sucrose 0.075M, free-fatty acid BSA (Sigma-Aldrich) 0.1%, HEPES 2 mM pH 7.4 in distilled water), centrifuged again at 10500g for 10 min and finally resuspended in 100 µl of isolation buffer 2. Mitochondrial protein concentrations were determined by Bradford assay. Mitochondrial proteins were diluted in a 37°C warm mitochondrial assay buffer (MgCl2 5 mM, D-Mannitol 220 mM, KH2PO4 10mM, EGTA 1 mM, free-fatty acid BSA 0.2%, HEPES 2 mM, sucrose 70 mM pH 7.0 in distilled water) and 1µg was loaded in a 96-well plate and centrifuged at 2000g for 20 min. After centrifugation 135 µl of mitochondrial assay buffer completed with 20mM succinate and 2µM rotenone were gently added to each well and the plate was warmed up to 37°C for 10 min. The plate was then loaded into a calibrated Seahorse XF96 Extracellular Flux Analyzer for mitochondrial respiration assay using the mito stress test kit (Seahorse Bioscience, # 103015-100). Mitochondrial respiration assay consisted in equilibration, 1 min mixing, 3 min pause, 1 min mixing, 3 min waiting, 0.5 min mix, 3 min measure, 1 min mixing, 3 min measure, 0.5 min mixing, injection of 4 mM ADP, 1 min mixing, 3 min measure, 1 min mixing, injection of 3.125 µM oligomycin, 0.5 min mixing, 3 min measure, 1 min mixing, injection of 4 µM FCCP, 0.5 min mixing, 3 min measure, 1 min mixing, injection of 4 µM antimycin, 0.5 min mixing, 3 min measure. All steps before plate loading were performed at 4°C.

### Mitochondrial calcium uptake assay

Quadriceps, tibialis anterior (TA) and EDL muscles of both legs from each mouse were pooled together, rinsed 2 times with PBS and minced finely with scissors in 0.5 ml of mitochondrial isotonic buffer (mannitol 225 mM, sucrose 75 mM, MOPS 5 mM, EGTA 0.5 mM, taurine 2 mM, pH 7.25). The resulting muscle homogenates were incubated 3 min in 10ml of mitochondrial isotonic buffer containing nargase at 0.1 mg/ml (Sigma-Aldrich, P8038) before the addition of BSA at 0.2%. Muscle solutions were then transferred in Potter-Elvehjem grinders and homogenized with manual pestles. Homogenates were centrifuged for 6 min at 1200g for 2 times, discarding fat and cellular debris between centrifugations. Obtained supernatants were centrifuged for 10 min at 9000g. Residual fat was discarded and mitochondrial pellets were washed with 15ml of mitochondrial isolation buffer before a second centrifugation at 9000g for 10 min. Final mitochondrial pellets were gently resuspended in a final volume of 150 µl of mitochondrial isotonic buffer. Mitochondrial protein concentrations were determined by Bradford method and 275 µg of mitochondria were centrifuged and gently resuspended in 100 µl of mitochondrial calcium assay buffer (KCl 120 mM, Tris 10 mM, MOPS 5 mM, K2HPO4 5 mM, pH 7.4). 100 µl of calcium buffer containing 5 µM of the fluorescent calcium indicator calcium green 5N (Molecular probes, C3737) was loaded in a 96-well black plate (Nunc) and baseline fluorescence was measured using a Tecan infinite M1000 multiplate reader. When the signal was stable, mitochondrial preparations were added and calcium green 5N fluorescence was continuously recorded during subsequent addition of calcium.

### FDB fiber isolation and calcium measurements

The flexor digitorum brevis (FDB) muscle was dissected manually from 28 months old male mice anaesthetized with isoflurane (4%) and killed by cervical dislocation. The FDB muscle was enzymatically dissociated for 1 hour in Tyrod’s buffer (138 mM NaCl, 2 mM CaCl2, 1mM Mg acetate, 4 mM KCl, 5 mM glucose, 10 mM HEPES, pH 7.4) containing 2.2 mg/ml collagenase I (Sigma-Aldrich) in the incubator at 37 °C and 5% CO2. After incubation with collagenase, muscle fibers were manually isolated using fire polished pipette tips and transferred onto matrigel coated 35mm glass bottom dishes (Ibidi GmbH, Martinsried, Germany). The muscle fibers were kept in Dulbecco’s Modified Eagle Medium (DMEM) supplemented with 10 % FCS and 1 % penicillin-streptomycin in the incubator at 37°C for 3-4 hours before medium was exchanged.

For the assessment of mitochondrial calcium uptake during electrical stimulation, FDB fibers were stained with Rhod-2 AM (ThermoFischer Scientific, Waltham, US) dissolved in Tyrod’s buffer for 20 min at room temperature. The final concentration of the dye in buffer was 1µM. During the calcium measurements N-benzyl-p-toluene sulphonamide (BTS, Sigma-Aldrich) was added to the buffer at a concentration of 10 µM to prevent muscle fiber contractions, without affecting calcium release.

Calcium imaging was carried out on an inverted Olympus FV3000-IX83 microscope equipped with a 20x UPSAPO lens. A uni-directional line scan mode with a temporal resolution of 0.488ms/line (2000 lines in total) was used to measure the temporal profile of the fluorescence signal. All experiments were carried out at room temperature, which was kept constant at 20 °C. Calcium release from the sarcoplasmic reticulum of FDB fibers was induced by a 1ms, 9V single electrical pulses using linear platinum electrodes that were placed at the edges of the glass bottom dish in about 1cm distance from each other. Typically, 50 fibers from one dish could be analyzed within a 20min time frame.

For image analysis we used Icy (open source software created by the Quantitative Image Analysis Unit at Institut Pasteur, Paris, France) in combination with the CalciumFluxAnalysis plugin to quantify the fluorescent intensity profiles in terms of amplitude (ΔF/F) and kinetics. The statistical analysis was carried out in Systat (Systat Software, Inc.).

### Transmission electron microscopy

Samples were prepared mitochondria density was calculated as previously described (73) and digitally adapted to Adobe Photoshop. Volume density is the ratio of test points residing within mitochondria and the total amount of test points within the field of view. Mitochondria were outlined manually. Using the measurement tool in Photoshop, the number of pixels contained in each mitochondrion were compared to the total number of pixels in the image. Mitochondrial size for was measured as the pixel area contained in each mitochondrion, adjusted to µm^2^ scale corresponding to image magnification. 5 images were quantified per block of stained muscle tissue, for a total of 5 blocks per mouse. Averages of mitochondrial size were taken across all mitochondria measured per mouse.

### Statistical analysis

Data were analyzed with two-way ANOVA (GraphPad Prism software). Sidak post-tests were used for multiple comparison analysis following two-way ANOVA. All data are plotted as mean ± S.E.M. For calcium measurements in FDB fibers, statistical tests were based on a mixed model analysis (SYSTAT 13) and the animal group was used as the fixed factor in each statistical analysis.

## Author Contributions

**Conceptualization:** J.F.G., J.D., S.M., J.S.P. C.H.

**Formal analysis:** J.F.G., J.D., S.M., J.S.P., M.R., C.H.

**Investigation:** J.F.G., G.S., S.F., S.M., M. R., S.S.

**Writing** J.F.G, J.D., C.H.

